# Evidence that the protein phosphatase activity of PTEN contributes to embryonic development and tumour suppression in mice

**DOI:** 10.64898/2026.03.14.711778

**Authors:** Priyanka Tibarewal, Laura Spinelli, Nisha Kriplani, Helen Wise, Nadege Poncet, Giulia Marzano, Karen E Anderson, Katarzyna M Grzes, Zofia Varyova, Mahreen Adil, C. Peter Downes, Phillip T Hawkins, Len R Stephens, Kate G Storey, Doreen A Cantrell, Bart Vanhaesebroeck, Nicholas R Leslie

## Abstract

PTEN (phosphatase and tensin homologue deleted on chromosome ten) is a tumour suppressor, the function of which is impaired in many diverse cancers. It has phosphoinositide lipid phosphatase activity by which it suppresses activation of the oncogenic PI3K signalling network but *in vitro* also displays activity against protein substrates and is able to auto-dephosphorylate its Thr366 residue. Here we generate germline knock-in mice expressing PTEN-Y138L, a mutant enzyme which selectively lacks protein phosphatase activity and retains lipid phosphatase activity. Homozygous *Pten*^*Y138L/Y138L*^ mice die *in utero* before E10.5. Primary MEFs and thymocytes with only a single *Pten*^*Y138L*^ allele display normal low levels of AKT phosphorylation indicating effective regulation of PI3K signalling by endogenous PTEN-Y138L *in vivo*. Heterozygous *Pten*^*+/Y138L*^ mice have reduced overall survival compared to wild type littermates and develop tumours in multiple organs. Our data imply that in addition to its lipid phosphatase activity, the protein phosphatase activity of PTEN is also required for normal embryonic development and tumour suppression.

## Introduction

Changes which reduce the function of the PTEN tumour suppressor are amongst the most commonly observed events in human sporadic tumours [1, 2]. And individuals with a germline *PTEN* mutation display diverse phenotypes including an increased risk of malignancy, leading to the designation PTEN Hamartoma Tumour Syndrome (PHTS) [3, 4]. PTEN is a member of the protein tyrosine phosphatase family which by dephosphorylation of PIP_3_ and PI(3,4)P_2_ executes an evolutionarily conserved role functionally opposing the activity of the class I PI 3-kinases (PI3Ks) and hence suppressing the activity of AKT, mTOR (mechanistic target of rapamycin) and other PI3K-dependent regulatory pathways [2, 5, 6]. PI3K signalling represents one of the most consistently activated functional pathways in cancer [7, 8]. Accordingly, mice heterozygous for a null allele of *Pten* and knock-in mice carrying a *Pten*^*G129E*^ mutation which selectively ablates the lipid phosphatase activity of PTEN both develop an array of tumours [9–12]. Remarkably, the tumour phenotype of *Pten*^*+/G129E*^ mice is slightly more severe than that of *Pten*^*+/−*^ animals [10, 12]. This implies not only that the lipid phosphatase activity is key to tumour suppression but also identifies a dominant negative effect mediated by lipid phosphatase-inactive PTEN. On the other hand, many other potential PI3K-independent mechanisms of action for PTEN have been identified, including phosphatase-independent functions in the nucleus [2, 13]. Notably, PTEN also displays weak but robust phosphatase activity *in vitro* against protein and phospho-peptide substrates, with highest activity against acidic phospho-tyrosine substrates [14], and a number of potential protein substrates have been proposed [15–20]. However, confident determination of the substrates of protein phosphatases is challenging and to date a clear picture is yet to emerge regarding the significance of these proposed substrates in PTEN function. PTEN also appears to slowly auto-dephosphorylate its C-terminal tail, specifically at Thr366 [21], a conclusion supported by structural data showing exposure of the PTEN active site upon dephosphorylation of Thr366 and Ser370 [22]. Importantly this latter study shows that phosphorylation of Ser370 and Thr366 strongly and selectively inhibit the activity of PTEN against soluble substrates which implies an evolved mechanism to control phosphatase activity against non-lipid substrates [22].

To test the significance of the protein phosphatase activity of PTEN, we previously engineered a PTEN mutant, PTEN-Y138L, which retains activity against lipid substrates yet lacks the normal activity of PTEN against phospho-peptides or Ins(1,3,4,5)P_4_, the soluble headgroup of PIP_3_ [23]. While *PTEN-Y138L* variants have not been reported in human disease, probably because mutation of Tyrosine 138 to Leucine involves two nucleotide changes, there are currently 16 reported cased of other mutations at *PTEN-Y138* in the COSMIC database (https://cancer.sanger.ac.uk/cosmic) mainly in the endometrium (7/16 cases), with 6 cases of *PTEN-Y138C*, 5 of *PTEN-Y138\**, 4 of *PTEN-Y138D* and one of *PTEN-Y138S*. PTEN-Y138C, like PTEN-Y138L, retains its PIP_3_ phosphatase activity but lacks protein phosphatase activity [21]. PTEN-Y138* leads to premature truncation of the PTEN protein and complete loss of expression and therefore function. The impact of Y138D and Y138S on PTEN function is not known. One PHTS patient has been reported with a *de novo PTEN-Y138C* variant [24].

We and others have since used PTEN-Y138L to study the role of PTEN’s protein phosphatase activity in regulating diverse cellular and physiological processes. Both PTEN-Y138L and PTEN-Y138C retain the ability to suppress AKT phosphorylation in cultured cells yet fail to inhibit glioma cell invasion [21] or control 3D lumen formation in mouse mammary epithelial NMuMG cells [25] unlike the wild-type PTEN enzyme. Notable in these studies, the activity of PTEN-Y138L in both cell-based assays could be rescued by mutation of Thr366 but did require lipid phosphatase activity. This indicates that, at least in these assays, the only requirement for the protein phosphatase activity of PTEN is the auto-dephosphorylation of this residue in the PTEN C-terminus [21, 25].

Additional studies have found that in human mammary epithelial MCF10A cells PTEN-Y138L has similar stability to wild-type PTEN but exhibits partial loss of function in spheroid formation assays [26]. Expression of GFP-PTEN-Y138L in *Pten*-null rat hippocampal sections fails to rescue increased neuronal spine density, implicating protein phosphatase activity in neuronal morphology. This activity appears to operate through auto-dephosphorylation of Ser380/Thr382/Thr383 residues [27]. Additionally PTEN-Y138L has also been linked to embryonic developmental defects in zebrafish models [28].

Some studies using PTEN-Y138L and the lipid phosphatase-dead PTEN-G129E mutant have shown that PTEN’s protein phosphatase activity plays a role in double stranded (ds) DNA damage repair and in DNA interstrand crosslink (ICL) repair, although the underlying mechanisms remain to be elucidated [29–31].

To further dissect the role of PTEN’s protein phosphatase activity *in vivo*, we generated knock-in mice expressing *Pten*^*Y138L*^. We show that homozygous *Pten*^*Y138L/Y138L*^ mice die *in utero* suggesting that PTEN’s protein phosphatase activity is essential for normal mouse embryonic development. Previously, we showed that breeding mice with only a single *Pten*^*Y138L*^ allele in the prostate does not induce tumorigenesis, initially suggesting that protein phosphatase activity is dispensable for PTEN’s tumour suppressor function in this organ. [32]. In contrast, our current findings of ubiquitous *Pten*^*Y138L*^ expression in heterozygous *Pten*^*+/Y138L*^ mice show tumour development across multiple organs, highlighting a requirement for PTEN’s protein phosphatase activity in tumour suppression.

## Results

### PTEN-Y138L inhibits AKT phosphorylation as efficiently as PTEN-WT in cultured cells

We have previously shown that expression of wild-type PTEN or PTEN-Y138L led to a similar reduction in AKT phosphorylation in the glioma cell line U87-MG, the triple negative breast cancer cell line MDA-MB-468 and the prostate cancer cell line LNCaP which all lack functional PTEN [21, 23, 25, 32].To exclude the possibility that this result was cell-type specific or an artefact of overexpression, we used lentiviral delivery to conduct experiments with a range of PTEN expression levels in U87-MG (**Fig. 1A**), MDA-MB-468 (**Fig. S1**) and in HCT116 colorectal cancer cells from which PTEN had been deleted (**Fig. 1B**) [33]. Increasing expression of PTEN caused a dose-dependent reduction in the phosphorylation of several proteins recognised as components or targets of the PI3K signalling network, including the downstream PIP3-regulated protein kinase AKT, the AKT substrates FOXO1 and PRAS40, as well as the mTOR substrate, ribosomal protein S6 (**Fig. 1 and S1**). In agreement with previous data [21, 23, 24], no differences were observed between the effects of PTEN wild-type and PTEN-Y138L expression on AKT phosphorylation or other components of the PI3K network (**Fig. 1 and S1**).

**Figure 1.**
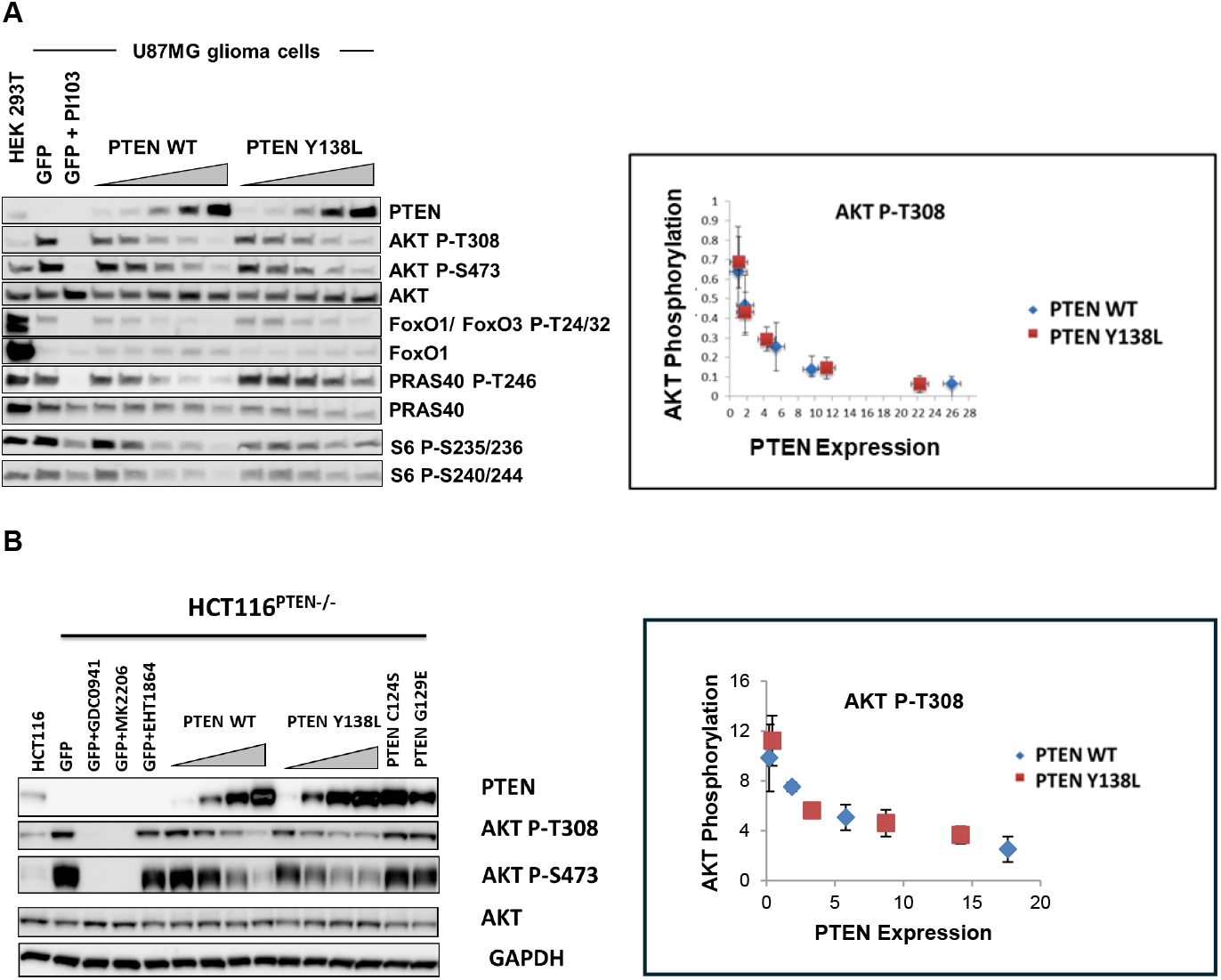
Characterisation of the effects of PTEN-Y138L on PI3K/AKT signalling: (**A**) U-87 MG and (**B**) isogenic HCT116 wild-type and *PTEN* knock out (−/−) cells transduced with fixed or increasing concentrations of lentiviruses for GFP, PTEN-WT or indicated PTEN mutants were either left untreated or treated as indicated with 1µM of PI103, GDC0941, MK2206 or EHT1864 for 1hr, followed by immunoblot analysis for the identified antibodies. Representative blots from n=3. The graphs show quantification of AKT P-308 at different concentrations of PTEN-WT and PTEN-Y138L.

### Homozygous PTEN-Y138L mice die *in utero*

To explore directly the correlation of protein phosphatase activity with the *in vivo* function of PTEN, we modified the endogenous *Pten* gene in mice by generating germline *Pten*^*Y138L*^ knock-in mutant mice (**Fig. S2**). We initially inter-crossed heterozygous *Pten*^*+/Y138L*^ mice. Among the 254 mice born, none were homozygous indicating that the PTEN-Y138L mutation leads to embryonic lethality in mice (**Fig. 2A**). To determine the stage of embryogenesis at which the homozygous embryos die, we examined embryos at E8.5 and found that homozygous embryos lacked anterior neural folds and appeared smaller in size than wild type and heterozygous embryos. At E9.5 the homozygous embryos had failed to undergo embryonic turning and were much smaller than wild type and heterozygous embryos and by E10.5 the homozygous embryos were reabsorbed by the uterine lining suggesting that homozygous embryos were no longer viable by this stage of embryonic development (**Fig. 2A**).

**Figure 2.**
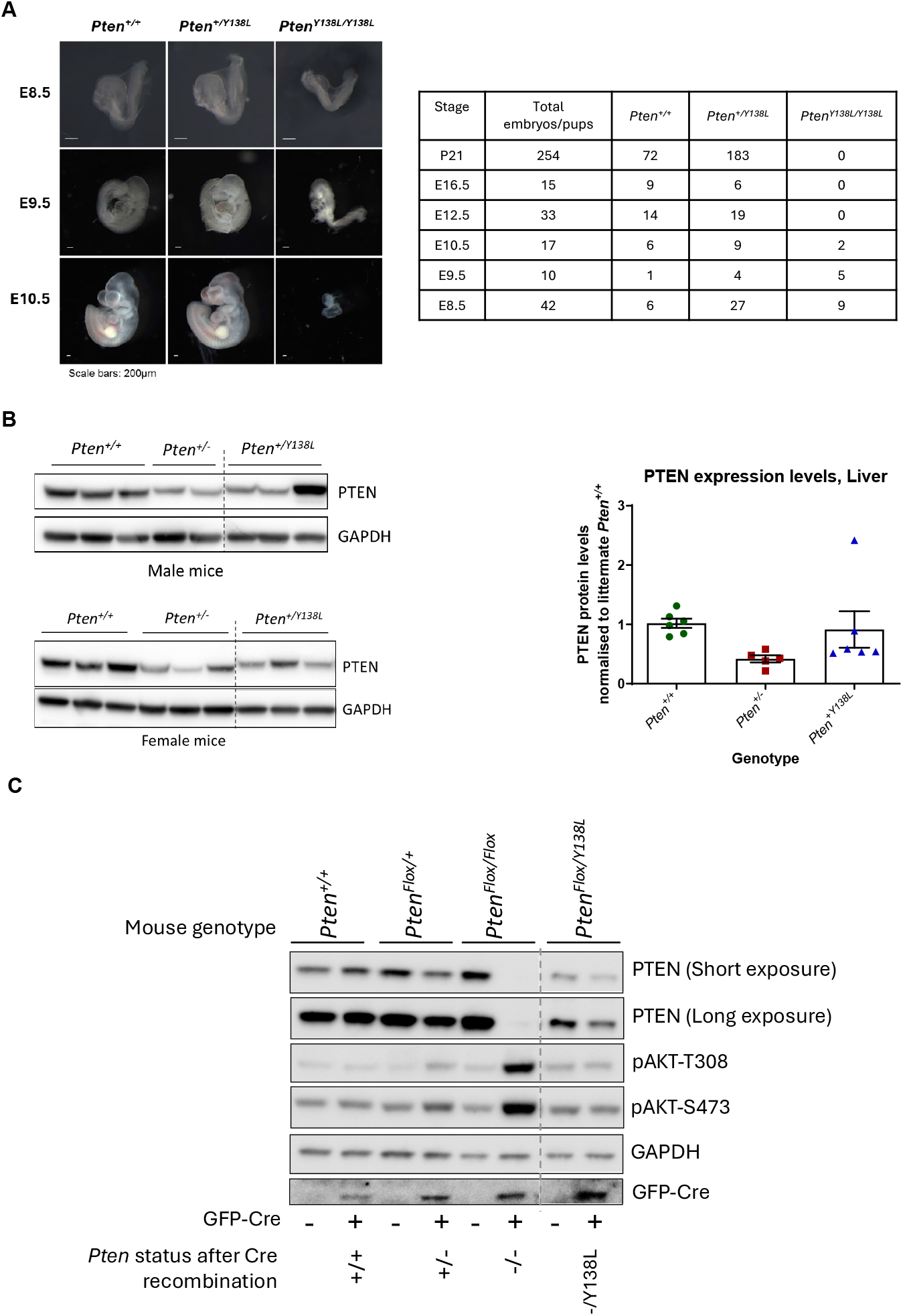
Characterisation of *Pten*^*Y138L*^ mice: (**A**) Homozygous *Pten*^Y138L/Y138L^ mice die *in utero* after embryonic stage E9.5. Images show morphology of E8.5, E9.5 and E10.5 *Pten*^+/+^, *Pten*^+/Y138L^ and *Pten*^Y138L/Y138L^ embryos. Scale bar: 200µm. Numerical data show the number of embryos/pups of each genotype obtained at the indicated time points. (**B**) Protein extracts from liver tissue of male (top left) and female mice (bottom left) of the indicated genotype were used for immunoblotting to assess PTEN expression. The graph on the right shows quantification of immunoblots. Data are shown as mean±SEM. PTEN expression normalised to GAPDH levels and relative to *Pten*^*+/+*^ littermate mouse. At least n=3 mice were used for each sex and each genotype except for male *Pten*^*+/−*^ where n=2 was used. (**C**) MEFs of the indicated genotype were transduced with GFP-Cre expressing lentiviruses. 96 hrs post transduction, cells were used for immunoblotting with the antibodies shown. Representative immune blots from n=3 experiments.

### Characterisation of *Pten*^*Y138L*^ mice

Since homozygous *Pten*^*Y138L*^ mice were not viable, we initially characterised heterozygous *Pten*^*+/Y138L*^ mice alongside age matched *Pten*^*+/+*^ as controls. These mice were on a C57BL/6J background. Mice heterozygous for *Pten (Pten*^*+/−*^) on a C57BL/6J have been previously characterised by us and other groups [9, 34–37] and were used as additional controls. In previous studies we found that PTEN-Y138L was less stable than wild-type PTEN, being expressed at 30-50% of the levels of PTEN-WT [23]. Cycloheximide chase experiments confirmed that PTEN-Y138L had a slightly shorter half-life compared to PTEN-WT **(Fig. S3A)**. Therefore, to ensure that any phenotypes attributable to the Pten-Y138L mutation are not simply due to lower levels of PTEN expressed in these mice and cells, we included a *Pten*^*hyp*^ hypomorphic allele in some of our experiments, which expresses approximately 50% reduced levels of the wild-type PTEN protein (**Fig. S3B** and [38]). Analysis of liver tissue from these mouse lines showed expression levels of PTEN were lower in tissues from heterozygous *Pten*^*+/−*^ mice than in wild-type mice and intermediate levels in *Pten*^*+/Y138L*^ mice (**Fig. 2B**).

We assess whether the PTEN-Y138L protein was expressed efficiently from the endogenous gene locus, we crossed *Pten*^*+/Y138L*^ mice with mice carrying a conditionally deletable *Pten*^*flox*^ allele to generate *Pten*^*flox/Y138L*^ mice. Immunoblot analysis of MEFs from these mice expressing Cre recombinase using retroviruses to knock out expression of the wild-type *Pten* allele, showed that PTEN protein is expressed from the mutant *Pten*^*Y138L*^ allele (**Fig. 2C**). Cre-expressing MEFs from *Pten*^*flox/flox*^ mice and which lack detectable PTEN protein have a substantially higher level of AKT phosphorylation relative to Cre-expressing MEFS from *Pten*^*flox/Y138L*^ mice confirming that the PTEN-Y138L protein retains its ability to inhibit PI3K-dependent signalling.

### Reduced overall survival of *Pten*^*+/Y138L*^ mice compared to wild-type mice

We next aged *Pten*^*+/Y138L*^ mice alongside *Pten*^*+/−*^, *Pten*^*+/Hyp*^ and *Pten*^*+/+*^ mice. Mice were monitored regularly and euthanised when presented with palpable lumps or signs of sickness, or at the age of 700 days. Consistent with what we have previously reported [34, 39], female *Pten*^*+/−*^ mice have reduced overall survival (median survival 199 days) compared to male mice (median survival 229 days). Both male and female *Pten*^*+/Y38L*^ mice had a significantly reduced overall survival when compared to wild-type *Pten*^*+/+*^ mice, with shorter survival in female *Pten*^*+/Y138L*^ mice relative to male *Pten*^*+/Y138L*^ mice (median survival 468 days vs 586 days respectively) (**Fig. 3A**). Overall survival of *Pten*^*+/Y138L*^ mice was however significantly higher than *Pten*^*+/−*^ mice (**Fig. 3A**). *Pten*^*+/Hyp*^ mice had a similar overall survival as wild-type *Pten*^*+/+*^ mice (**Fig. 3A**).

**Figure 3.**
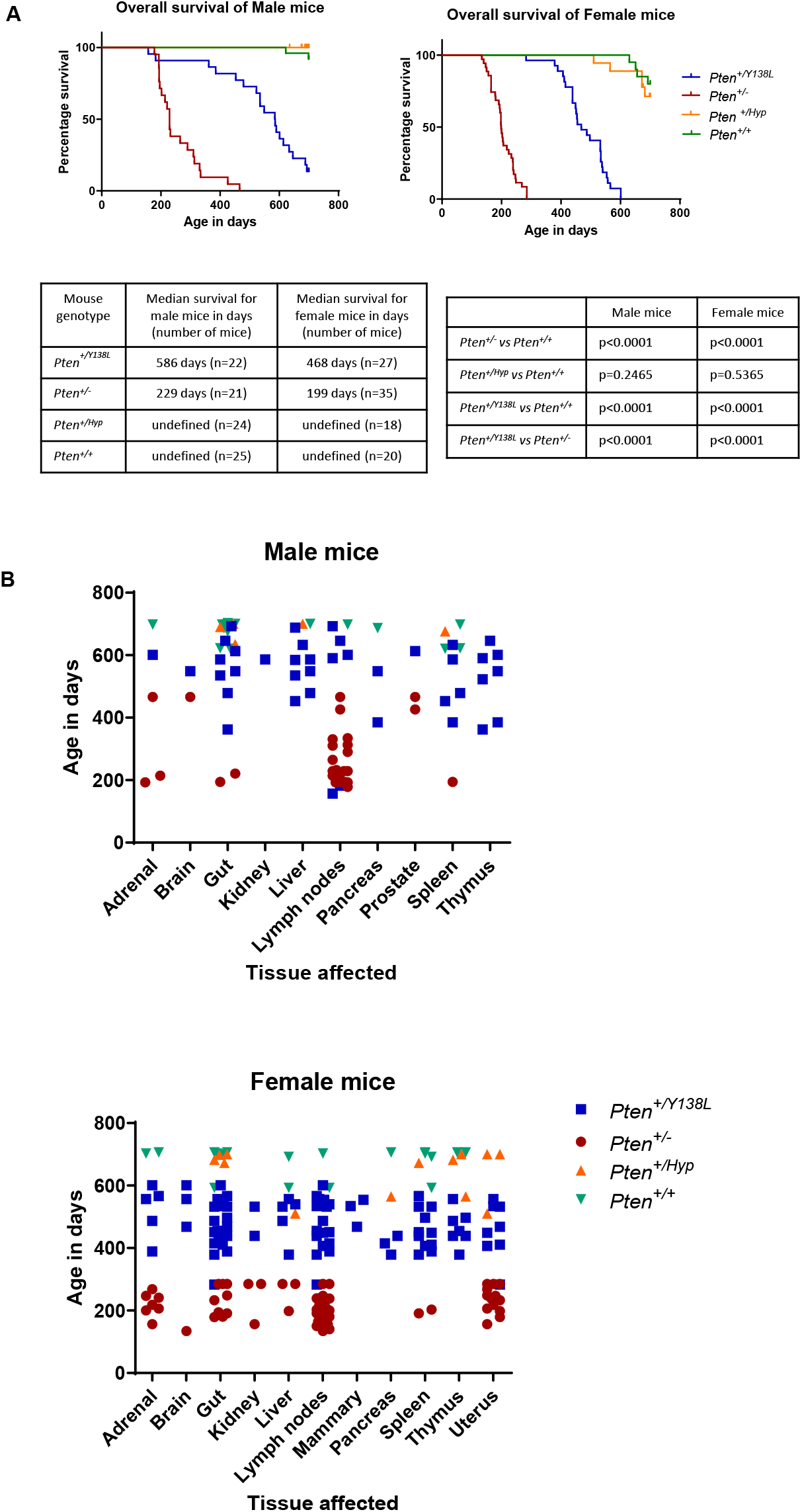
Survival and histopathological analysis of *Pten*^+/Y138L^ mice. *Pten*^+/Y138L^, *Pten*^*+/Hyp*^ and *Pten*^+/−^ mice were allowed to age alongside wild-type *Pten*^*+/+*^ littermates. Mice were euthanized for welfare reasons (ill health or palpable masses) or at a specified age (730 days). (**A**) Kaplan–Meier survival curves for male and female mice. Numerical data show the median survival age of the mice. Pairwise comparisons were made between genotypes and statistical analysis was performed using the Log-rank (Mantel–Cox) test and p values are shown to the right. **(B)** Incidence of lesions in the indicated tissues in male (top) and female (bottom) mice, as assessed by histopathological analysis on all mice, showing the age of lesion identification and sacrifice (Y-axis) and the mouse genotype (colour coded following the key shown).

### *Pten*^*+/Y138L*^ mice develop tumours in multiple organs

Mice heterozygous for null alleles of *Pten* have been studied extensively as they spontaneously develop a range of tumour types, the spectrum of which appears to be influenced strongly by the genetic background of the mice [9]. We and others have previously reported that *Pten*^*+/−*^ mice on a C57BL/6J background almost all develop B-cell lymphomas before 12 months of age [9, 35, 40–42]. Other tumour types included carcinoma of the prostate, endometrium and mammary tissue, and adrenal pheochromocytoma. To study spontaneous tumour development in *Pten*^*+/Y138L*^ mice and compare it to that in *Pten*^*+/−*^, *Pten*^*+/Hyp*^ and *Pten*^*+/+*^ mice, we performed post-mortem analysis of mice that were euthanised as part of the aging study. A summary of our findings is shown in Table 1. Figure 3B shows the age distribution of the individual mice with lesions in the indicated tissue.

**Table 1:**
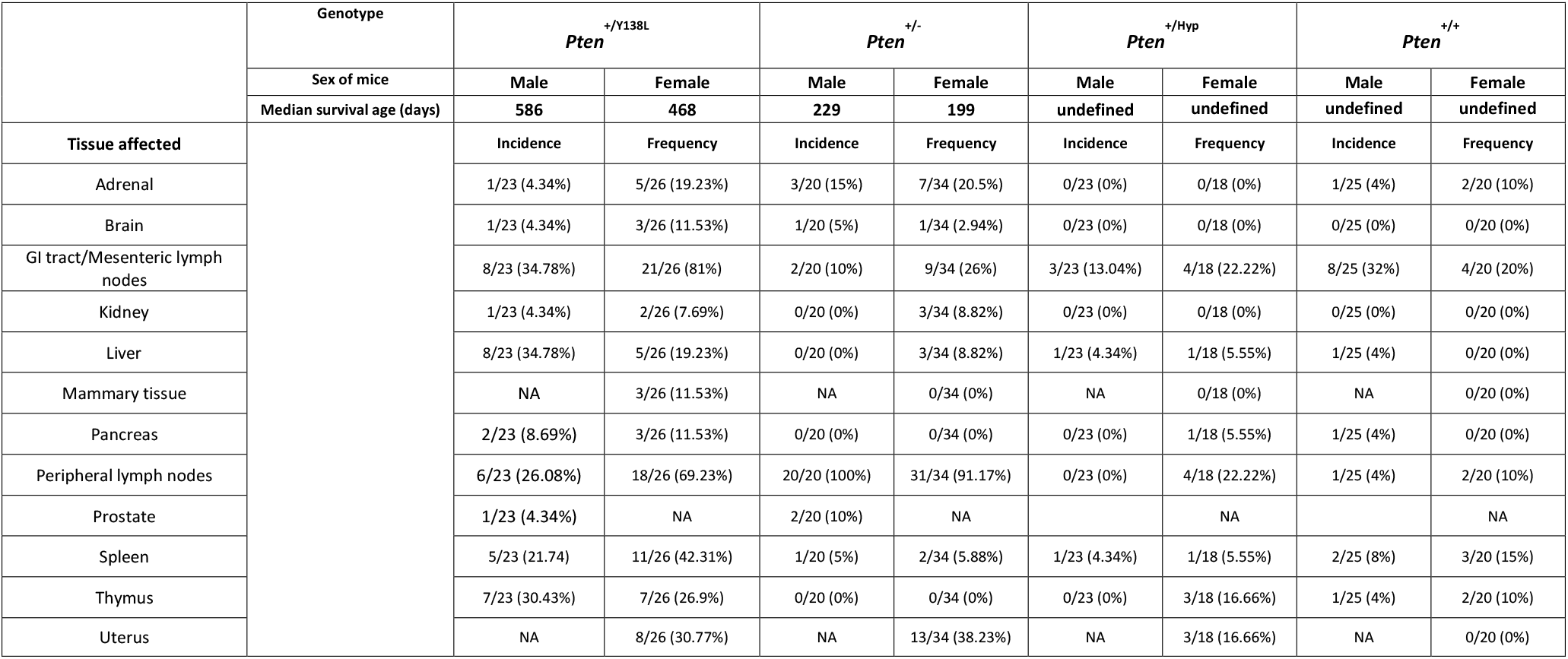
Incidence and frequency of lesions found in *Pten* mutant and wild type mice.

|  | Genotype | <sup>+ / Y138L</sup><br>*Pten* |  | <sup>+ / -</sup><br>*Pten* |  | <sup>+ / Hyp</sup><br>*Pten* |  | <sup>+ / +</sup><br>*Pten* |  |
| --- | --- | --- | --- | --- | --- | --- | --- | --- | --- |
|  | Sex of mice | Male | Female | Male | Female | Male | Female | Male | Female |
|  | Median survival age (days) | 586 | 468 | 229 | 199 | undefined | undefined | undefined | undefined |
| Tissue affected |  | Incidence | Frequency | Incidence | Frequency | Incidence | Frequency | Incidence | Frequency |
| Adrenal |  | 1/23 (4.34%) | 5/26 (19.23%) | 3/20 (15%) | 7/34 (20.5%) | 0/23 (0%) | 0/18 (0%) | 1/25 (4%) | 2/20 (10%) |
| Brain |  | 1/23 (4.34%) | 3/26 (11.53%) | 1/20 (5%) | 1/34 (2.94%) | 0/23 (0%) | 0/18 (0%) | 0/25 (0%) | 0/20 (0%) |
| GI tract/Mesenteric lymph nodes |  | 8/23 (34.78%) | 21/26 (81%) | 2/20 (10%) | 9/34 (26%) | 3/23 (13.04%) | 4/18 (22.22%) | 8/25 (32%) | 4/20 (20%) |
| Kidney |  | 1/23 (4.34%) | 2/26 (7.69%) | 0/20 (0%) | 3/34 (8.82%) | 0/23 (0%) | 0/18 (0%) | 0/25 (0%) | 0/20 (0%) |
| Liver |  | 8/23 (34.78%) | 5/26 (19.23%) | 0/20 (0%) | 3/34 (8.82%) | 1/23 (4.34%) | 1/18 (5.55%) | 1/25 (4%) | 0/20 (0%) |
| Mammary tissue |  | NA | 3/26 (11.53%) | NA | 0/34 (0%) | NA | 0/18 (0%) | NA | 0/20 (0%) |
| Pancreas |  | 2/23 (8.69%) | 3/26 (11.53%) | 0/20 (0%) | 0/34 (0%) | 0/23 (0%) | 1/18 (5.55%) | 1/25 (4%) | 0/20 (0%) |
| Peripheral lymph nodes |  | 6/23 (26.08%) | 18/26 (69.23%) | 20/20 (100%) | 31/34 (91.17%) | 0/23 (0%) | 4/18 (22.22%) | 1/25 (4%) | 2/20 (10%) |
| Prostate |  | 1/23 (4.34%) | NA | 2/20 (10%) | NA |  | NA |  | NA |
| Spleen |  | 5/23 (21.74) | 11/26 (42.31%) | 1/20 (5%) | 2/34 (5.88%) | 1/23 (4.34%) | 1/18 (5.55%) | 2/25 (8%) | 3/20 (15%) |
| Thymus |  | 7/23 (30.43%) | 7/26 (26.9%) | 0/20 (0%) | 0/34 (0%) | 0/23 (0%) | 3/18 (16.66%) | 1/25 (4%) | 2/20 (10%) |
| Uterus |  | NA | 8/26 (30.77%) | NA | 13/34 (38.23%) | NA | 3/18 (16.66%) | NA | 0/20 (0%) |

Macroscopic observations showed that the majority of *Pten*^*+/Y138L*^ mice had masses in, or attached to, the gut (21/26 female mice and 8/23 male mice), at higher frequency than in *Pten*^*+/−*^ mice (9/34 female mice and 8/23 male mice) (**Fig.3B and Table 1**). Histological analysis of the GI tract from a fraction of these mice revealed that the masses attached to the GI tract were large B-cell lymphomas of the mesenteric lymph nodes. The masses inside the GI tract were juvenile polyps which were histopathologically similar to those seen on *Pten*^*+/−*^ mice. A significant number of mice had enlargement of peripheral lymph nodes (18/26 female mice and 6/23 male mice) (**Fig.3B and Table 1**). Histologically, these were B-cell lymphomas similar to what has been reported previously in *Pten*^*+/−*^ mice and were also seen in all our *Pten*^*+/−*^ mice (20/20) and was the most common cause of euthanasia (**Fig. 3B and Table 1**). Female *Pten*^*+/Y138L*^ mice developed endometrial hyperplasia (8/26 mice) and mammary carcinoma (3/26), while 1/23 male *Pten*^*+/Y138L*^ mouse developed prostate carcinoma (**Fig.3B and Table 1**). Both *Pten*^*+/Y138L*^ and *Pten*^*+/−*^ mice also developed adrenal pheochromocytomas (**Fig.3B and Table 1)**. Although there is no reported link between PTEN and pathogenesis of human pheochromocytomas, it is commonly observed in *Pten* mouse models [43].

Age and genetic background related tumours were seen in *Pten*^*+/Y138L*^, *Pten*^*+/Hyp*^ and *Pten*^*+/+*^ mice and included lymphomas of the liver, pancreas, spleen and thymus (**Fig.3B and Table 1**). These were either absent or seen at a low frequency in *Pten*^*+/−*^ mice probably because these mice were euthanised at a young age because of the earlier development of other tumours and did not develop tumours that occur later in life in a PTEN mutant background (**Fig.3B and Table 1**).

### Loss of PTEN’s protein phosphatase activity does not lead to development of T-cell leukemias

PTEN function is frequently lost in human T-cell leukemias and mice lacking PTEN in T-lymphocytes develop T-cell lymphomas in young adulthood [44, 45]. Using mouse models expressing *Pten*^*C124S*^ and *Pten*^*G129E*^, it has been shown that loss of PIP_3_ phosphatase activity of PTEN is key the development of T-cell lymphomas [46]. To investigate the role of the PTEN protein phosphatase activity in T-cell development and T-cell lymphomagenesis, we intercrossed *Pten*^*+/Y138L*^ mice, *Pten*^*Flox*^ mice and mice which express *Lck-Cre* to delete the *Pten*^*Flox*^ allele specifically in T-cells. This generated genotypes including *Pten*^*Flox/Y138L*^ *Lck-Cre*, heterozygously expressing PTEN-Y138L ubiquitously, with their T-cells expressing only PTEN-Y138L from the endogenous allele, alongside *Pten*^*Flox/Flox*^ *Lck-Cre* which completely lack PTEN expression in the thymocytes but are otherwise wild-type for PTEN. Mice with wild type thymocytes (*Pten*^*+/+*^) and thymocytes expressing only one copy of *Pten (Pten*^*+/Flox*^ *Lck-Cre*) were used as additional controls.

Western blot analysis on thymocytes of 5-week-old mice showed that as expected the expression levels of PTEN in *Pten*^*Flox/Y138L*^ *Lck-Cre* thymocytes was less compared to *Pten*^*Flox/+*^ *Lck-Cre* and *Pten* ^*+/+*^ wild-type thymocytes (**Fig. 4A**). However, the phosphorylation of AKT was not increased in either *Pten*^*Flox/Y138L*^ *Lck-Cre* or *Pten*^*Flox/+*^ *Lck-Cre* thymocytes but was elevated in *Pten*^*Flox/Flox*^ *Lck-Cre* thymocytes (**Fig. 4A**). Analysis of lipid extracts from primary thymocytes from these mice showed that *Pten*^*Flox/Flox*^ *Lck-Cre* mice have significantly elevated levels of PIP_3_ when compared to *Pten*^*+/+*^ mice, whereas thymocytes from *Pten*^*Flox/Y138L*^ *Lck-Cre* and *Pten*^*Flox/+*^ *Lck-Cre* mice did not display this increase (**Fig. 4B**).

**Figure 4.**
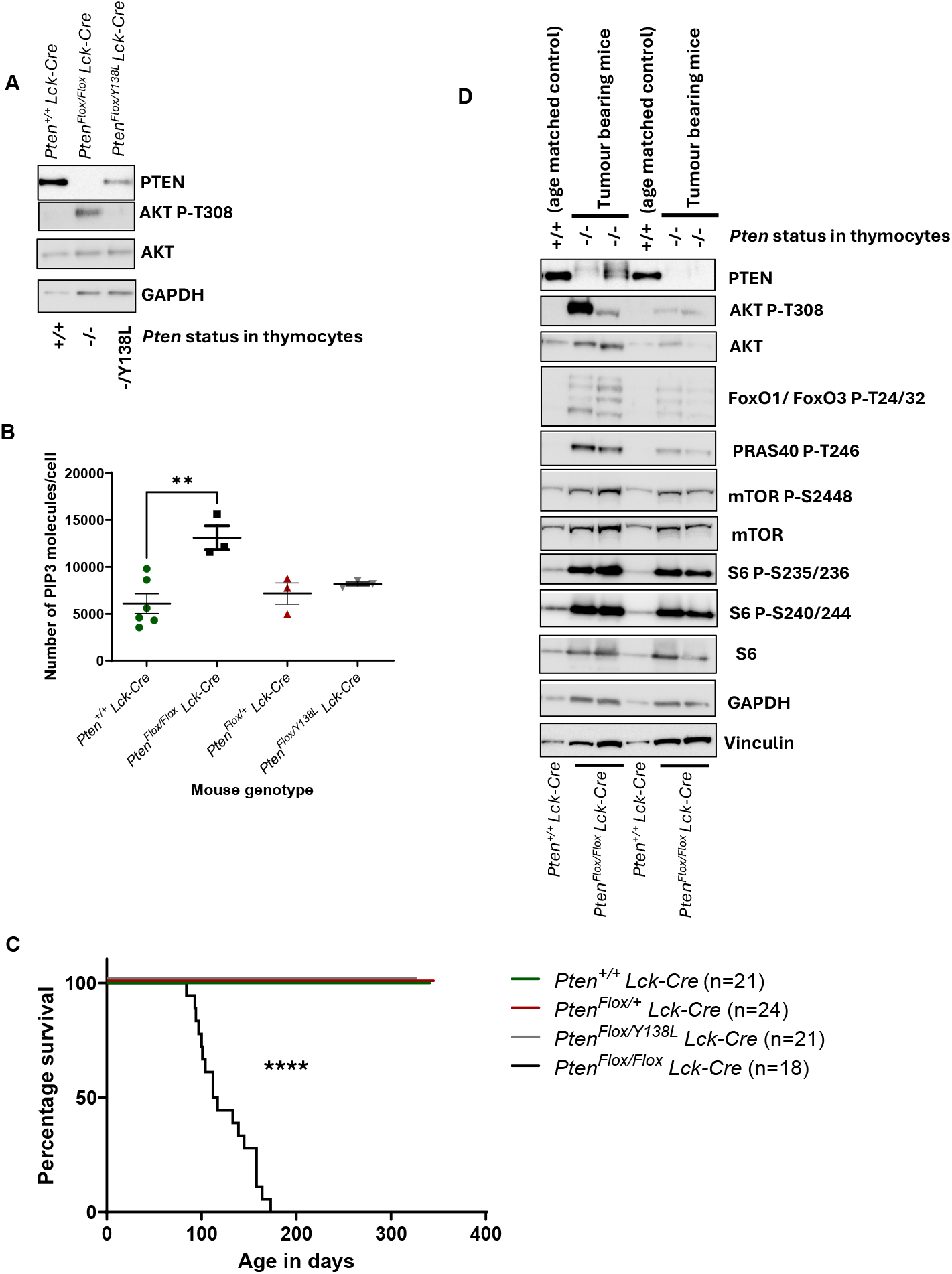
Analysis of mice with monoallelic expression of *Pten*^*Y138L*^ in the T-cell lineage: (**A**) Protein extracts from thymocytes isolated from 5-week-old *Pten*^*Flox/Y138L*^ *Lck-Cre, Pten*^*Flox/Flox*^ *Lck-Cre* and *Pten*^*+/+*^ *Lck-Cre* mice were used for immunoblotting for the indicated antibodies. A representative blot from n=3 experiments is shown. (**B**) Thymocytes isolated from 5-week-old *Pten*^*Flox/Y138L*^ *Lck-Cre, Pten*^*Flox/Flox*^ *Lck-Cre, Pten*^*Flox/+*^ *Lck-Cre* and *Pten*^*+/+*^ *Lck-Cre* mice were used for PIP_3_ measurements using Mass Spectrometry. The graph shows mean±SEM from n=3 mice for all genotypes except *Pten*^*+/+*^*Lck-Cre* for which n=6 was used. Statistical analysis was performed using One Way ANOVA (ANOVA p=0.0050; **) followed by Tukeys post hoc test (*Pten*^*+/+*^ *Lck-Cre* vs *Pten*^*Flox/Flox*^ *Lck-Cre* p=0.0031; **). (**C**) *Pten*^*Flox/Y138L*^ *Lck-Cre, Pten*^*Flox/Flox*^ *Lck-Cre, Pten*^*Flox/+*^ *Lck-Cre* and *Pten*^*+/+*^ *Lck-Cre* mice were allowed to age alongside wild-type *Pten*^*+/+*^ littermates. Mice were euthanized for welfare reasons (ill health or palpable masses) or at a specified age (300 days) and their survival presented as Kaplan– Meier plots. Pairwise comparisons were made to *Pten*^*+/+*^*Lck-Cre* mice and statistical analysis was performed using Log-rank (Mantel–Cox) test (*Pten*^*+/+*^*Lck-Cre* vs *Pten*^*Flox/Flox*^ *Lck-Cre* p value<0.0001; ****). (**D**) Protein extracts of thymocytes from tumour bearing *Pten*^*Flox/Flox*^ *Lck-Cre* (n=4) and age matched control *Pten*^*+/+*^*Lck-Cre* (n=2) mice were used for immunoblot analysis of for the indicated antibodies.

We allowed a cohort of *Pten*^*Flox/Y138L*^ *Lck-Cre* mice to age alongside *Pten*^*Flox/+*^ *Lck-Cre* mice, *Pten*^*Flox/Flox*^ *Lck-Cre* and *Pten* ^*+/+*^ wild-type mice to study development of spontaneous T-cell lymphomas (**Fig. 4C**). Consistent with previous reports, *Pten*^*Flox/Flox*^ *Lck-Cre* mice developed aggressive T-cell lymphomas with a median survival age of about 16 weeks (114.5 days) (**Fig. 4C**). These lymphomas characterised by the monoclonal expansion of T-cells had increased levels of phosphorylation of AKT and other downstream effectors of the PI3K/AKT signalling pathway such as Foxo1/3, mTOR and S6 ribosomal protein (**Fig. 4D** and [44, 47]). In contrast, *Pten*^*Flox/Y138L*^ *Lck-Cre* mice did not develop T-cell lymphomas during the 10 month course of this study (cohort culled at the age of 300 days) (**Fig. 4C**). Consistent with the near normal PIP_3_ levels, *Pten*^*Flox/+*^ *Lck-Cre* mice did not develop T-cell lymphomas. These results suggest T-cell lymphoma initiation and progression depends on the increase of PIP_3_ levels and PI3K-AKT signalling and therefore is regulated by the PIP_3_ phosphatase function of PTEN.

## Discussion

The tumour suppressor role of PTEN has been mainly attributed to its activity as a lipid phosphatase. Mouse models carrying null or hypomorphic *Pten* alleles, or the catalytically inactive mutants *Pten*^C124S^ or the selectively lipid phosphatase-deficient *Pten*^*G129E*^, develop tumours across multiple organs, overlapping with the tumour spectrum in PHTS patients [9, 10, 12, 42, 48]. Evidence from other mouse models has highlighted the role of lipid phosphatase-independent functions of PTEN in tumour suppression. Notably, mice expressing a nuclear-excluded mutant *Pten*^*R173C*^, which retains the ability to regulate the canonical PI3K/AKT signalling pathway also develop tumours, although with delayed onset, most likely driven by genomic instability caused by impaired double stranded (ds) DNA damage repair [39]. Likewise, mice heterozygously expressing PTEN which lacks the C-terminal PDZ-binding sequence show accelerated development of mammary tumours induced by polyoma middle T-antigen or MYC, suggesting that lack of interaction with PDZ domain containing proteins plays a role in the development of these tumours [49, 50].

Here we characterise a mouse model expressing a protein phosphatase-deficient *PTEN* mutant *Pten*^*Y138L*^, which retains its lipid phosphatase activity. Our results show that homozygous *Pten*^*Y138L/Y138L*^ animals cease developing in utero around embryonic day 8.5. The smaller size of these embryos indicates a broad impact on development potentially due to impacts on placenta development [51]. Heterozygous *Pten*^*+/Y138L*^ mice also have a reduced overall survival compared to wild-type littermates and develop a similar spectrum of tumours, albeit delayed, as observed in *Pten*^*+/−*^ mice.

Our data therefore correlate the protein phosphatase activity of PTEN with normal embryonic development and tumour suppression. The data are consistent with previous experiments in cultured cells which indicate that the lipid phosphatase activity of PTEN alone provides a strong suppression of AKT and proliferation, but that both protein and lipid phosphatase activities must act together to control the more complex biological processes of glioma cell invasion and epithelial tissue architecture [21, 23, 25]. Co-expression of PTEN-Y138L and PTEN-G129E did not suppress glioma cell invasion or support normal epithelial morphogenesis, implying that both activities must exist within the same PTEN molecule, perhaps to allow regulation by auto-dephosphorylation [21, 25]. However, it is not clear whether these cultured cell-based assays represent *bona fide* models of tumour suppression *in vivo*. Therefore, it is also possible that full tumour suppressor activity *in vivo* requires PTEN to dephosphorylate both PIP_3_ and one or more heterologous protein substrates.

The PTEN-Y138L mutant protein is somewhat less stable than wild-type PTEN, raising the possibility that this reduced stability and resultant lower protein concentration leads to reduced PIP_3_ metabolism and tumour suppressor function. To control for this, we included in our analysis of spontaneous tumour formation mice heterozygous for a hypomorphic allele of *Pten* and observed a much weaker tumour phenotype in these mice than in our *Pten*^*+/Y138L*^ mice. It is also noteworthy that the expression of PTEN-Y138L alone from the endogenous gene at slightly lower levels than PTEN-WT is sufficient to suppress AKT phosphorylation in MEFs and thymocytes. Additionally, in a quantitative yeast-based functional analysis of >7000 (86% of all possible) PTEN missense mutants, PTEN-Y138L was found to retain strong activity, consistent with its ability to metabolise cellular PIP_3_ [52]. These findings argue against a protein-dose dependent effect caused by unstable PTEN-Y138L being a dominant factor in driving the formation of tumours and embryonic lethality in mice expressing PTEN-Y138L. Instead, they argue strongly for the functional importance of a qualitative functional deficit which correlates with the loss of protein phosphatase activity. One plausible explanation is that PTEN’s protein phosphatase activity is required for effective dsDNA damage repair, and partial impairment of this function in *Pten*^*+/Y138L*^ mice may predispose to tumour development [30]. Or that the lipid and protein phosphatase activities of PTEN are required in concert to coordinate metabolism of small, localised pools of PIP_3_. However, we cannot exclude the alternative possibility that cell-type specific differences in PTEN-Y138L stability and degradation prevent its accumulation and function in tumour-initiating cells, despite its apparent activity in MEFs and thymocytes.

We observed that a single copy of either *Pten*^*Y138L*^ or wild-type *Pten* was sufficient to suppress the formation of T-cell lymphomas caused in mice by the deletion of both *Pten* gene copies in T-cells. This shows that the suppression of these murine T-cell lymphomas correlates with the ability of PTEN to suppress canonical PI3K-AKT signalling and is also in agreement with similar observations in the murine prostate [32].

Despite extensive research, spatiotemporal dynamics of PTEN function and regulation remain poorly understood. Ongoing investigations are addressing these gaps, particularly in the context of therapeutic strategies targeting PTEN-deficient cancers, such as the use of PI3K-AKT pathway inhibitors. [53–55]. Our findings, alongside those of others, underscore the clinical relevance of these regulatory nuances. They highlight the potential for mechanistic insights into PTEN’s biochemical functions to inform the rational use of existing therapies and inspire novel treatment approaches that more effectively reflect PTEN’s diverse tumour suppressive roles.

## Acknowledgements

The authors thank Stewart Fleming for analysis of mouse pathology. We also thank Dario Alessi and Lloyd Trotman for sharing genetically modified mice and Todd Waldman for sharing PTEN null HCT116 cells. We would also like to thank the staff of the Biological Resource Unit at the University of Dundee for help with in vivo work. This work was supported by a grant from the UK Medical Research Council (Ref: G0801865).

## Methods

### Cell lines

U-87 MG and MDA-MB-468 cells were purchased from ECACC, and cultured in Minimum Essential Medium (MEM) (Gibco) supplemented with 10% fetal bovine serum (FBS) (Gibco). HEK293T cells were purchased from ECACC and Lenti-X™ 293T were purchased from Clontech, Phoenix cells were purchased from ATCC, and cultured in Dulbecco’s Modified Eagle Medium (DMEM) with 10% FBS. Isogenic HCT-116 wildtype and *PTEN* knock out cells were a gift from Todd Waldman (Georgetown University) and were cultured McCoy’s modified medium with10% FBS. All cells were grown under standard cell culture conditions at 37°C in a 5% CO_2_ humidified incubator.

### Plasmids

Generation of lentiviral constructs for pHRSIN-PTEN-WT, Y138L, C124S, G129E have been described before [23]. pBabe-Hygro-HTert and pHRSIN-GFP-Cre plasmids were obtained from the Division of Signal Transduction Therapy, Dundee, UK.

### Lentiviruses and retroviruses

Lentiviral particles encoding PTEN wild-type, PTEN mutants, GFP or GFP-Cre were generated by co-transfecting Lenti-X™ 293T or HEK-293T cells with plasmids containing the relevant cDNA and packaging vectors pHR-CMV 8.2 deltaR and pCMV VSV-G, using TransIT-LT1 (Mirus Bio) according to the manufacturer’s protocol. Retroviral particles were produced by transfecting Phoenix cells with pBabe-Hygro-Htert using the same transfection reagent and protocol. 24 h hours post-transfection, sodium butyrate was added to a final concentration of 12.5 mM for 6 hours, followed by PBS washes and replacement with fresh media. Lentiviral supernatants were harvested after 20 hours, filtered through a 0.45 µm membrane, and stored at −80°C. For transduction, target cells were seeded at 40–50% confluency. After cell attachment, lentiviral or retroviral particles were added along with polybrene (Sigma-Aldrich) at 20 µg/µl. Media was refreshed 24 h post-transduction.

### Cycloheximide chase studies

U-87 MG cells were transduced with lentiviral constructs encoding PTEN-WT or PTEN-Y138L. 48 h post-transduction, cells were treated with cycloheximide at a final concentration of 200 µg/ml. At designated time points following treatment, cells were lysed, and protein extracts were collected for immunoblot analysis of PTEN abundance.

### Immunoblotting analysis

Protein extracts from cultured cells were obtained by scraping cells into ice-cold lysis buffer containing: 25 mM Tris-HCl (pH 7.4), 150 mM NaCl, 1% Triton X-100, 10% glycerol, 1 mM EGTA, 1 mM EDTA, 5 mM sodium pyrophosphate, 10 mM β-glycerophosphate, 50 mM sodium fluoride, 1 mM sodium orthovanadate, 1 mM DTT, and a protease inhibitor cocktail (Millipore). Mouse tissues collected post-euthanasia were homogenized in Lysing Matrix M tubes (MP Biomedicals) using twice the tissue volume (v/w) of lysis buffer composed of: 25 mM Tris-HCl (pH 7.4), 150 mM NaCl, 1% Triton X-100, 0.1% SDS, 10% glycerol, 1 mM EGTA, 1 mM EDTA, 10 mM sodium pyrophosphate, 20 mM β-glycerophosphate, 100 mM sodium fluoride, 2 mM sodium orthovanadate, 1 mM DTT, and protease inhibitors. Homogenization was performed on a FastPrep-24 system (MP Biomedicals) at 4 m/sec for 20 sec. Lysates from cells and tissues were clarified by centrifugation at 20,000 × g for 10 min at 4°C, and the resulting protein extracts were subjected to immunoblotting.

For gel electrophoresis, protein extracts diluted with 4X LDS sample buffer (Thermo Fisher Scientific) were loaded onto NuPAGE Bis-Tris 4–12% gradient polyacrylamide gels (Thermo Fisher Scientific) and processed according to the manufacturer’s instructions. Proteins were transferred to PVDF membranes (Millipore), which were then blocked in 5% milk powder in TBST for 1 hour at room temperature. Membranes were incubated overnight with primary antibodies (Table S1), followed by a 1-hour incubation at room temperature with HRP-conjugated secondary antibodies (GE Healthcare). Signal detection was performed using Immobilon Forte Western HRP substrate (Millipore), and chemiluminescence was captured using the ImageQuant LAS4000 imaging system (GE Healthcare).

### Mice

All mice were maintained at University of Dundee in accordance with the UK Animals (Scientific Procedures) Act 1986 and following UK Home Office guidance. All procedures were authorized by a UK Home Office Project Licence (PPL 70/8128) subject to local ethical review. *Pten*^+/−^ mice [41], *Pten*^*+/Hyp*^ [38], *Pten*^*Flox/Flox*^ mice [56] and *Lck-Cre* mice [57] have been described elsewhere. *Pten*^*+/Y138L*^ mice were generated by Taconic Biosciences (Formerly Taconic Artemis, Cologne, Germany). Briefly a targeting vector was generated using BAC clones from the C57BL/6J RPCIB-731 BAC library containing a ~10kb region of *Pten* with exon 5 containing the Y138L mutation (c.412CA>T, c.413T>G (p.Tyr138Leu)), a puromycin resistant gene flanked by FRT recombinase sites in intron 4. A schematic representation of the targeting strategy is shown in **Fig.S2**. The targeting vector was then introduced in TaconicArtemis C57BL/6N Tac embryonic stem (ES) cell line by electroporation. Homologous recombinant clones were isolated using positive (PuroR) and negative (thymidine kinase-TK) selection. They were then screened by PCR and southern blot to confirm correct integration. Verified ES cell clones were injected into blastocysts, which were then implanted into pseudopregnant females to produce chimeric offspring. Chimeras were bred with wild-type mice to assess germline transmission of the modified *Pten* allele. Heterozygous progeny were genotyped and backcrossed with C57BL/6j mice to establish stable lines for downstream phenotypic and functional studies. All mice used were maintained on a C57BL/6j background.

### Genotyping mice

For genotyping, un-purified DNA was released from ear punch biopsys of mice taken at weaning or embryonic yolk sac by adding 20 μl of MicroLysis Plus (Microzone). The samples were placed in a Thermo Cycler used following the lysis program (65°C for 15 mins, 96°C for 2 mins, 65°C for 4 mins 96°C for 1 min, 65°C for 1 min, 96°C for 30 sec, 20°C hold). Released DNA was then amplified in PCR using the KAPA Biosystems 2G Fast HS PCR KIT (Sigma Aldrich) according to manufacturer’s instructions using primers listed in Table S2. The products of the PCRs were detected with gel electrophoresis.

### Mouse survival studies

All sample size calculations were performed with a Type I error probability (α) of 0.05 and a statistical power of 80%. For spontaneous tumour development studies, sample size estimates for *Pten*^+/Y138L^ mice were based on reported median survival ages of *Pten*^*+/−*^ mice on a C57BL/6J background (approximately 8–12 months [35, 37]). To detect a 20% improvement in median survival in *Pten*^+/Y138L^ mice relative to *Pten*^*+/−*^ controls, a minimum of 16 mice per sex was calculated. Accordingly, at least 16 mice of each sex were used for all genotypes studied in parallel: *Pten*^*+/Y138L*^, *Pten*^*+/+*^, *Pten*^+/−^ and *Pten*^*+/Hyp*^.

For T-cell lymphoma studies involving Pten *Pten*^*Flox/Y138L*^ *Lck-Cre* mice, sample size calculations were based on a reported median survival of 90–100 days for *Pten*^*Flox/Flox*^ *Lck-Cre* mice [44, 58]. To detect a 15% improvement in survival, a minimum of 8 mice per sex was calculated. At least 8 mice of each sex were included for all genotypes studied in parallel: *Pten*^*Flox/Y138L*^ *Lck-Cre, Pten*^*Flox/Flox*^ *Lck-Cre, Pten*^*Flox/+*^ *Lck-Cre* and *Pten*^*+/+*^ *Lck-Cre*.

For survival analysis, mice were monitored daily and euthanized upon exhibiting signs of ill health, including hunched posture, laboured breathing, lethargy, or palpable masses. Mice that remained asymptomatic were euthanized at 730 days (spontaneous tumour study) or 300 days (T-cell lymphoma study).

### Sequencing of *Pten* from mice

Liver tissue collected from mice were used for bulk RNA extraction using the RNeasy Mini Kit (Qiagen), according to the manufacturer’s instructions. To assess RNA integrity, 500 ng of RNA was electrophoresed on a 2% agarose gel, confirming the presence of distinct 28S and 18S rRNA bands at approximately 4.5 kb and 1.9 kb, respectively. cDNA was synthesized using the Sprint RT-random hexamer kit (Clontech), following the manufacturer’s protocol. Full-length PTEN cDNA was amplified by PCR using the primer pair listed in Table S3. PCR products were sequenced by the Sequencing Service at the University of Dundee, UK.

### Mouse embryo microscopy

Whole embryo images were taken using a Micropublisher 3.3RT and Q‐Imaging on a Leica MZFLIII dissecting microscope.

### Preparation of Mouse embryonic fibroblasts (MEFs)

To generate E13.5 mouse embryonic fibroblasts (MEFs), timed matings were performed between *Pten*^+*/Y138L*^ and *Pten*^*+/Flox*^ mice to generate MEFs used in **Fig. 2C** and between *Pten*^*+/+*^ and *Pten*^*+/Hyp*^ mice to generate MEFs used in **Fig. S3B**. Pregnant females were euthanized at embryonic day 13.5 (E13.5), and embryos were harvested from the uterus. After removal of the head and visceral organs, the remaining embryonic tissue was finely minced in 2× Trypsin/EDTA using a sterile scalpel and incubated at 37°C for 40 minutes. The resulting cell suspension was plated onto 10 cm dishes containing DMEM (Gibco) supplemented with 10% FBS (Gibco) and Penicillin-Streptomycin (Sigma-Aldrich) and cultured under standard conditions.

### Tissue processing for histological analysis

Immediately upon dissection, mouse tissues were fixed in 10% neutral buffered formalin for 24 hours, then processed and embedded into paraffin blocks. To ensure optimal sectioning, tissues were oriented with the largest surface facing down. Serial sections of 3 µm thickness were cut and stained with Haematoxylin and Eosin (H&E). All slides were evaluated by Prof Stewart Fleming.

### Thymocyte isolation

For isolation of thymocytes, mice were euthanised and the thymus was removed. They were then mashed with a syringe plunger in RPMI containing 10% FBS and filtered through a 70 µm filter. Cell suspension was centrifuged at 500xg for 5 mins to obtain a cell pellet which was washed once with PBS and recentrifuged. The cell pellets were then snap frozen in liquid nitrogen for immunoblot analysis of measurements of PIP_3_ levels.

### PIP_3_ measurements

PIP_3_ concentrations were measured from thymocytes using mass spectrometry (MS) as described previously [59, 60]. Briefly, 2.5×10^6^ thymocytes were used for lipid extraction followed by addition of 10ng of C17:0-C16:0 PIP_3_ as an internal standard. The HPLC-MS peak area of the internal standard served as a reference to calculate the absolute amount of PIP_3_ per cell, based on the corresponding sample peak areas. Cellular PIP_3_ appeared to be C38:4 PIP_3_.

### Statistical analysis

GraphPad Prism (San Diego, CA, United States) was used for statistical analysis. The method used for individual datasets is indicated in the figure legends.

