## Supplementary figures and tables for "Evidence that the protein phosphatase activity of PTEN contributes to embryonic development and tumour suppression in mice"

**This pdf consists of**

**Supplementary figures S1-S3**

**Supplementary tables 1-3**

**Fig. S1**

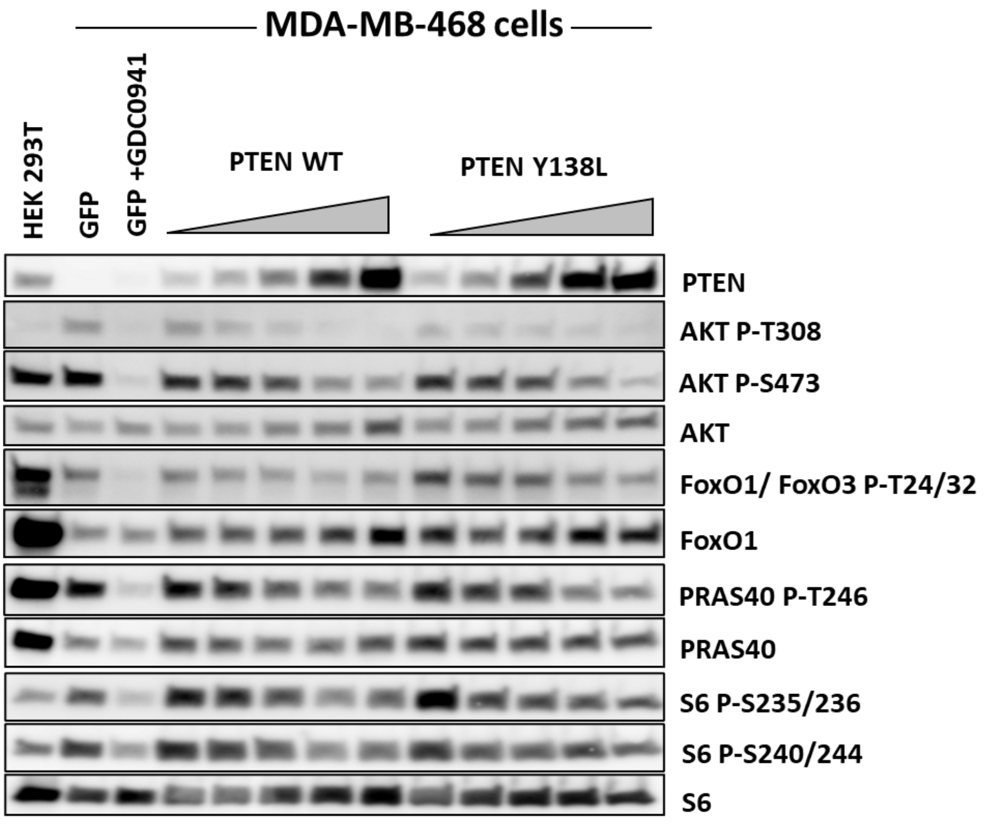


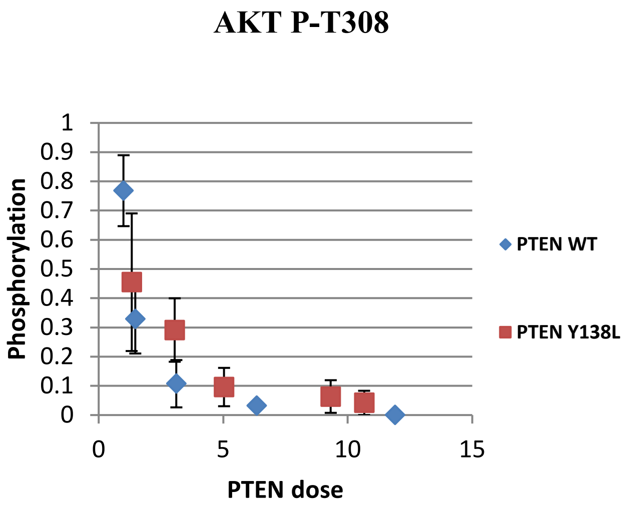


**Figure S1. Characterisation of the effects of PTEN-Y138L on PI3K/AKT signalling:** MDA-MB-468 cells transduced with fixed or increasing concentrations of lentiviruses for GFP, PTEN-WT or PTEN-Y138L were either left untreated or treated as indicated with 1µM of GDC0941 for 1hr, followed by immunoblot analysis for the identified antibodies. Representative blots from n=3. The graphs show quantification of AKT P-308 at different concentrations of PTEN-WT and PTEN-Y138L.

**Fig. S2**

**A**

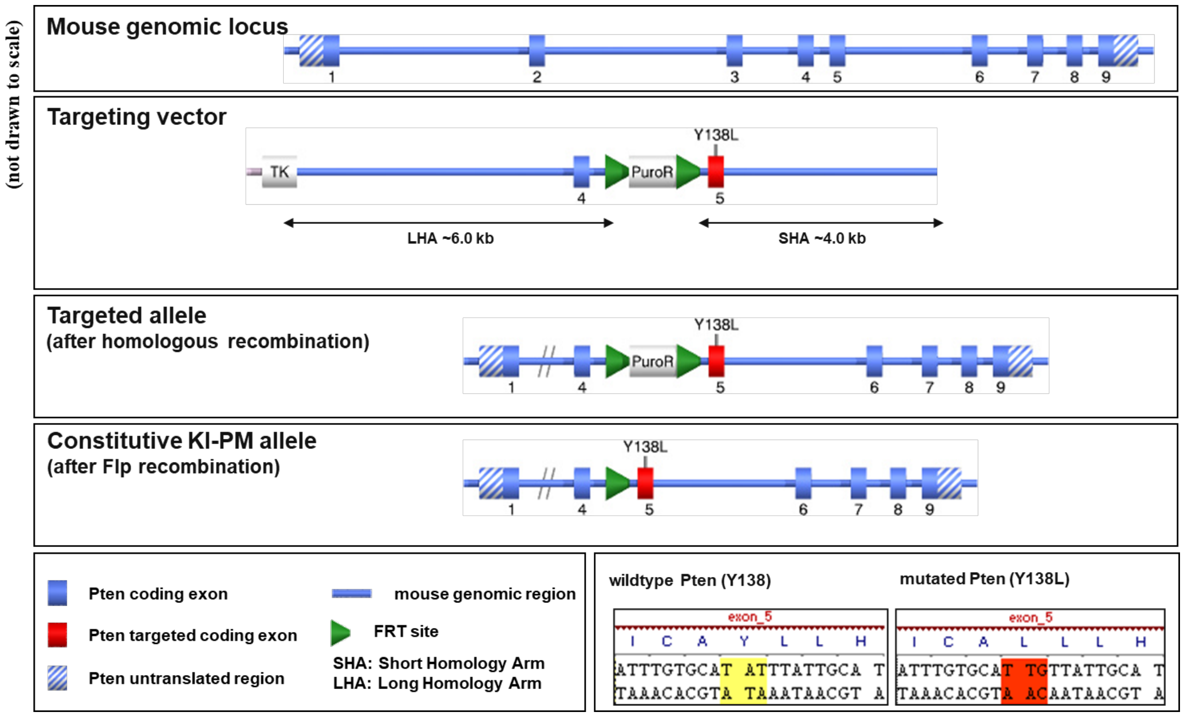


**B**

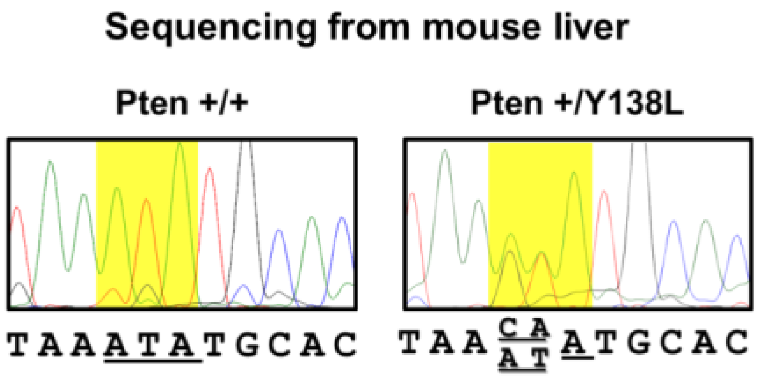


**Figure S2. Generation of *Pten*$^{Y138L}$ mice:** (**A**) Gene targeting strategy for generation of *Pten*$^{Y138L}$. Briefly a targeting vector was generated containing a ~10kb region of *Pten* with exon 5 containing the Y138L mutation (c.412CA>T, c.413T>G (p.Tyr138Leu)), a puromycin resistant gene flanked by FRT recombinase sites in intron 4. The targeting vector was then introduced into embryonic stem (ES) cell line by electroporation. Homologous recombinant clones were isolated and then implanted into pseudopregnant females to produce chimeric offspring, which were then bred with C57BL/6j mice to establish stable lines for further studies. (**B**) Sequence trace (3'-5' strand) from cDNA isolated from liver tissue of 8-week-old *Pten*$^{+/Y138L}$ and littermate *Pten*$^{+/+}$ mice showing heterozygous Y138L mutation.

**Fig. S3**

**A**

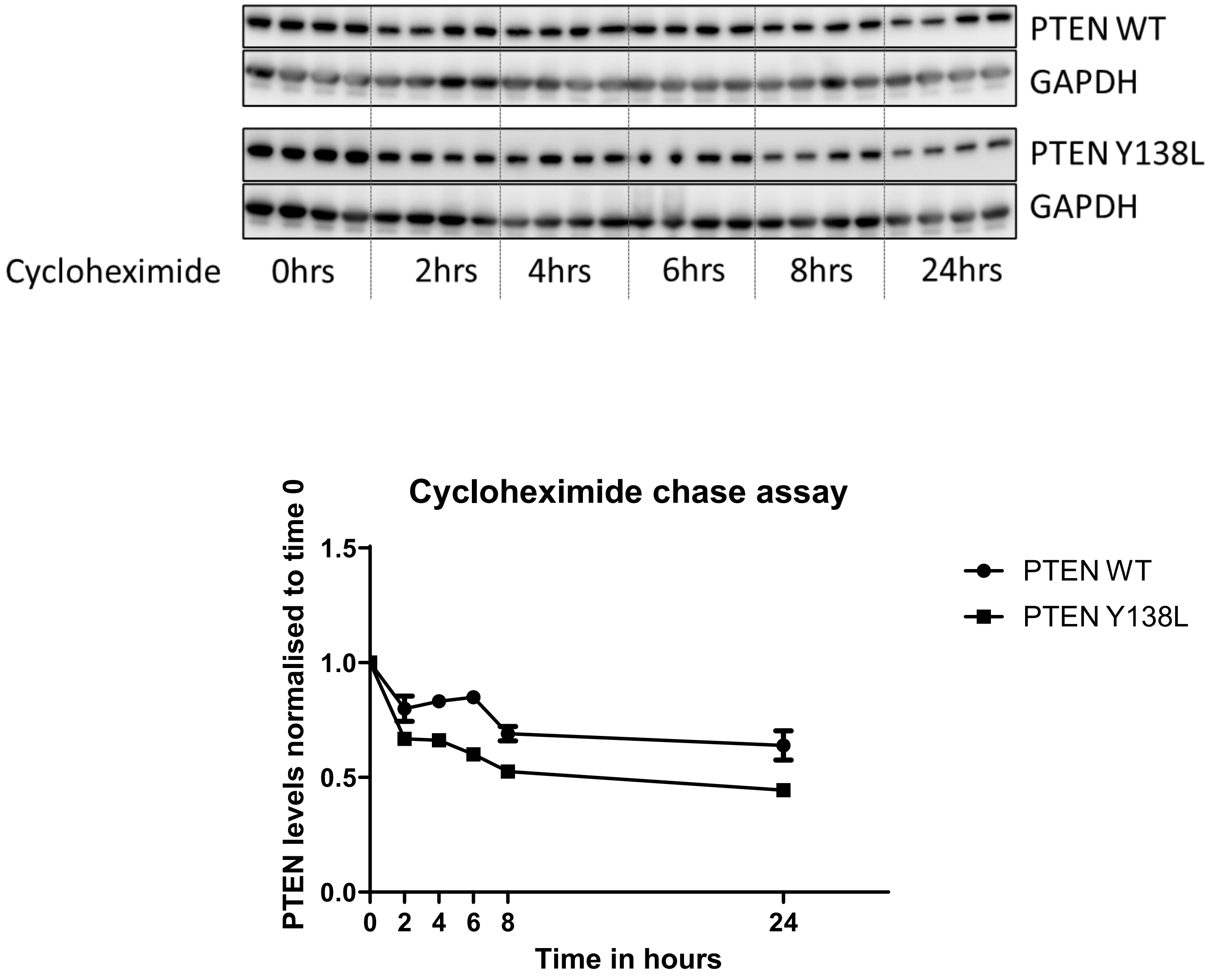


**B**

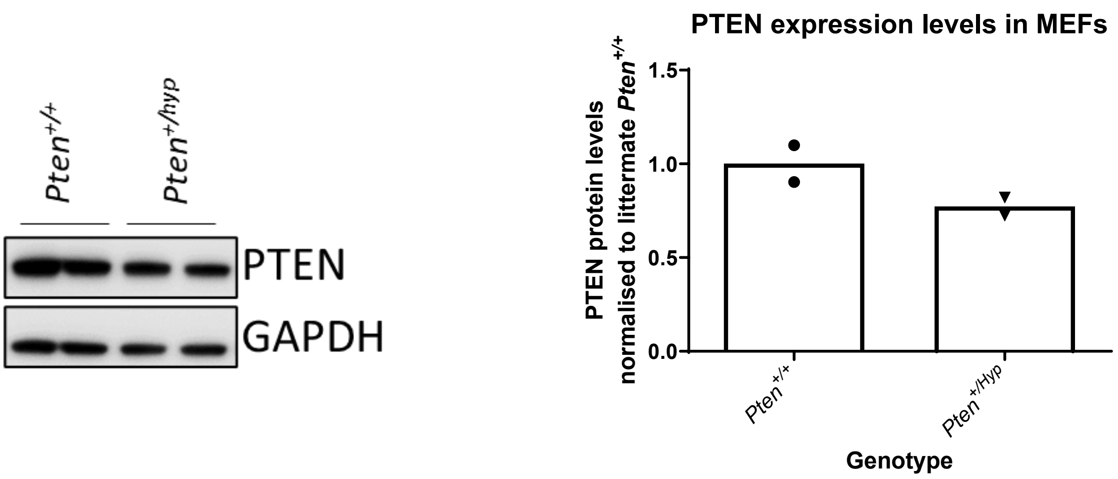


**Figure S3: Cycloheximide chase analysis of PTEN-Y138L and PTEN expression in MEFs:** (**A**) Transient lentiviral expression of PTEN wild-type (WT) or PTEN-Y138L in U-87 MG cells. The cells were treated with cycloheximide to inhibit protein synthesis, and PTEN protein levels were determined by immunoblotting at the indicated time points after cycloheximide treatment. The graph shows PTEN protein levels normalized to GAPDH levels relative to time 0, data shown as mean±SEM, n=4. (**B**) MEFs of the indicated genotype were used for immunoblotting with the antibodies shown. The graph on the right shows quantification of immunoblots. PTEN expression normalised to GAPDH levels and relative to *Pten*$^{+/+}$ littermate mouse. Data are shown as mean from n=2.

List of supplemetary tables

**Table S1: Antibodies used for western blotting**
**Table S2: Primers used for genotyping mice**
**Table S3: Primers for sequencing *Pten***

**Table S1: Antibodies used for western blotting**

| Antibody | Supplier | Catalogue number | Species reactivity | Dilution |
|---|---|---|---|---|
| PTEN | Santa Cruz Biotechnology | Sc-7974 | Mouse | 1:1000 |
| PTEN | Cell Signaling Technology | 9552 | Rabbit | 1:1000 |
| AKT P-S473 | Cell Signaling Technology | 9271 | Rabbit | 1:1000 |
| AKT P-T308 | Cell Signaling Technology | 9275 | Rabbit | 1:500 |
| AKT | Cell Signaling Technology | 9272 | Rabbit | 1:1000 |
| S6 P-S240/244 | Cell Signaling Technology | 2215 | Rabbit | 1:1000 |
| S6 P-S235/236 | Cell Signaling Technology | 2211 | Rabbit | 1:1000 |
| S6 | Cell Signaling Technology | 2217 | Rabbit | 1:1000 |
| PRAS40 P-T246 | Cell Signaling Technology | 2640 | Rabbit | 1:1000 |
| PRAS40 | Cell Signaling Technology | 2610 | Rabbit | 1:1000 |
| FoxO1 P-T24/FoxO3a P-T32 | Cell Signaling Technology | 9464 | Rabbit | 1:1000 |
| FoxO1 | Cell Signaling Technology | 2880 | Rabbit | 1:1000 |
| GAPDH | Cell Signaling Technology | 2218 | Rabbit | 1:10,000 |

**Table S2: Primers used for genotyping mice**

| Mouse line | Primers for genotyping | Expected size of PCR product |
|---|---|---|
| ***Pten*** $^{+/-}$ | *Pten* common: 5’ TTGCACAGTATCCTTTTGAAG 3’ | Wild-type: 240bp |
| | *Pten* WT: 5’ GTCTCTGGTCCTTACTTCC 3’ | Mutant: 320bp |
| | *Pten* Neo: 5’ ACGAGACTAGTGAGACGTGC 3’ | |
| ***Pten*** $^{+/Y138L}$ | PTEN-Y138L F: 5'-ATGGAAAGGAGTAAATGGATGG-3' | Wild-type: 250bp |
| | PTEN Y138L R: 5'-GGAGTAAAAGCAGGAGAATTGG-3' | Mutant: 300bp |
| ***Pten*** $^{+/Hyp}$ | PTEN 5’ Hypomorph: 5'-TGTTTTTGACCAATTAAAGTAGGCTGTG-3' | Wild-type: 350bp |
| | PTEN 3’ Hypomorph: 5'-AAAAGTTCCCCTGCTGATGATTTGT-3' | Mutant: 490bp |
| ***Pten*** $^{flox/flox}$ | *Pten* Flox F: 5’-GGCAAAGAATCTTGGTGTTAC-3’ | Wild-type: 230bp |
| | *Pten* Flox R: 5’-GCCTTACCTAGTAAAGCAAG-3' | *Pten* $^{flox}$ : 280bp |
| ***Lck-Cre*** | *Lck-Cre* F: 5' CGGTCGATGCAACGAGTGATGAGG 3' | PCR product of transgene at 600bp |
| | *Lck-Cre* R: 5' CCAGAGACGGAAATCCATCGCTCG 3' | |

**Table S3: Primers for sequencing *Pten***

PTEN 5’ UTR mouse F : 5'-CATGTTGCAGCAATTCACTG-3'
PTEN 3’ UTR mouse R: 5'-GGTATTTTATCCCTCTTGATAAG-3'
